# Trust-Aware Sequence-to-Function Modelling in Regulatory Genomics

**DOI:** 10.64898/2026.08.20.745945

**Authors:** Abdulmujeeb T. Onawole, Sulaimon Basiru, Musafau Oloyede Sanni, Monsuru Aiyedun, Ridwan Sulaimon

## Abstract

**Objective:** Sequence-to-function models increasingly predict regulatory activity, such as chromatin accessibility, directly from DNA sequence, and are used to interpret non-coding genetic variation. Standard accuracy metrics, computed over a held-out set of genomic regions, do not establish whether an individual prediction remains reliable once the input sequence departs from that set, nor whether a model’s attribution-based explanation is biologically grounded rather than coincidental. We develop and evaluate RegTrust-XAI, a trust-aware framework separating these questions using three inference-time signals: ensemble consensus, motif-grounded attribution coherence, and applicability-domain distance.

**Methods:** A five-model convolutional ensemble was trained on 517,790 K562 ATAC-seq windows and evaluated on a held-out chromosome test set (chr8/chr9, n = 42,844). Consensus, coherence, and applicability-domain distance were each tested against prediction error, alongside complementary sequence-novelty analyses and validation against an independent lentiMPRA reporter assay and saturation-mutagenesis MPRA data at the PKLR promoter.

**Results:** The ensemble reached Spearman *ρ* = 0.782, with skill of 0.328 over a constant-value null predictor. High-consensus predictions (Scenarios A+B) were consistently enriched for lower error than low-consensus predictions (Scenarios C+D), and attribution coherence further separated error within the high-consensus population (mean absolute error 0.396 versus 0.435, p = 9.6e-10). Applicability-domain distance showed a monotonic error gradient across six distance bands. A 4-mer composition-divergence metric was negatively associated with error and anti-correlated with applicability-domain distance, so composition-based and model-relevant novelty are not equivalent. Attribution transfer to lentiMPRA was assay- and subgroup-dependent, and predicted allele-substitution effects correlated with measured saturation-mutagenesis effects at the PKLR promoter at both 24 h and 48 h (*ρ* = 0.227 and 0.235). Motif-specific perturbation further showed that regulatory attributions were strongly context-dependent, with more than 90% of multi-instance motif modules exhibiting superadditive joint effects.

**Conclusions:** Prediction reliability, explanation validity, and sequence novelty are related but distinct properties of a sequence-to-function model. Evaluating each explicitly gives a more complete basis for deciding when to act on a prediction than accuracy alone.

## 1. Introduction

DNA regulatory sequence determines when and where a gene is transcribed, and models that predict regulatory activity directly from sequence are increasingly used to interpret non-coding genetic variation, prioritize candidate disease variants, and guide the design of synthetic regulatory elements. These sequence-to-function models take a DNA window as input and output a molecular readout, such as chromatin accessibility, transcription-factor occupancy, or reporter expression, learned from genome-wide assays such as ATAC-seq or massively parallel reporter assays, with the most recent models predicting several such readouts jointly from a single architecture [1]. The architecture used to make this prediction, whether a convolutional network [2, 3], a dilated convolutional network [4], or a transformer [5, 6], is not the question this study addresses. Recent architectural comparisons on this task class rarely produce a decisive winner once models are reasonably well tuned [7, 8].

A model that performs well on held-out genomic regions is not necessarily reliable for every new sequence. Existing genomes sample only a tiny and biased fraction of the possible regulatory-sequence space, so strong performance on a random or chromosome-based split does not guarantee similar performance on sequence variants, engineered constructs, or unfamiliar regulatory contexts [7]. Consistent with this, genomic deep-learning models can lose accuracy in cell-type-specific accessible regions despite strong genome-wide performance [9]. A recent personal-genome study provides an even clearer example, performance remained strong for unseen individuals at previously observed genes but collapsed for unseen alleles at unseen genes [10]. Importantly, that failure became apparent only because matched ground-truth expression data were available. This study therefore asks whether such failures can be flagged before ground truth is known using signals available at inference time.

A second, related problem concerns whether a model’s explanation of its own prediction can be trusted. Attribution methods highlight which input bases most influenced a prediction [11, 12], and when those bases overlap a known transcription-factor motif, the explanation looks biologically plausible. A plausible-looking explanation is not the same as a causally correct one. A documented case from a different domain shows an explainability method flagging sugar-ring substructures as the basis for a bitterness prediction, when the true cause was a dataset artefact in which natural products are frequently both glycosylated and bitter rather than glycosylation itself driving the taste [13, 14]. The equivalent risk in regulatory genomics is that a model’s attribution concentrates on a real transcription-factor motif for a reason that is statistically convenient rather than mechanistically load-bearing, so a correct prediction and a biologically grounded explanation need to be evaluated as separate claims rather than assumed to travel together.

This study develops and evaluates RegTrust-XAI, a trust-aware framework for assessing individual sequence-to-function predictions at inference time. The framework considers whether independently trained models agree on a prediction, whether the corresponding attribution is supported by relevant regulatory motifs, and whether the input lies within the model’s learned applicability domain. A related framework was previously developed for protein property prediction using ensemble consensus, explanation-based evidence, and applicability-domain distance [15]. Here, the same principle is adapted to regulatory genomics by grounding explanation coherence in cell-type-relevant transcription-factor motifs and assessing model familiarity in learned sequence space. We use K562 as the test system because it provides a well-characterized human model of regulatory biology with unusually rich functional-genomics data. K562 was derived from a patient with chronic myelogenous leukemia [16] and has been widely used to study hematopoietic and erythroid regulation [17]. It was also selected as an ENCODE Tier 1 cell line, making extensive chromatin, transcription-factor, and other regulatory datasets available in a common cellular context [18]. This combination makes K562 suitable for testing whether trust signals derived from a sequence-to-function model remain informative across held-out genomic regions, sequence perturbations, and independent regulatory assays. We therefore ask whether prediction stability, explanation validity, and model familiarity provide distinct information about the reliability of an individual regulatory prediction.

### Statement of Significance

| Summary | Description |
| --- | --- |
| **Problem or issue** | Sequence-to-function models are increasingly used to prioritize regulatory variants for translational and experimental follow-up, but standard accuracy metrics do not show whether any single prediction is reliable or whether its explanation reflects real regulatory biology. Without a signal for this at the point of prediction, there is no principled way to prioritize which results merit downstream follow-up and which need costly experimental confirmation first. |
| **What is already known** | Uncertainty estimation, explainability methods, and distribution-shift evaluation each address one facet of model trustworthiness in isolation. Existing reviews |
|  | identify calibrated uncertainty and biologically grounded explanation as open needs for sequence-to-function models but do not build or empirically test a working solution [7]. |
| **What this paper adds** | RegTrust-XAI combines three inference-time signals, ensemble consensus, motif-grounded attribution coherence, and applicability-domain distance, into one decision framework, validated against chromosome-held-out data, an independent reporter assay, and true saturation-mutagenesis measurements. It goes beyond prior conceptual reviews by empirically showing which signal combinations flag a prediction as a higher-confidence candidate for downstream prioritization versus one that still needs experimental confirmation. |
| **Who would benefit from the new knowledge in this paper** | Researchers and translational scientists who use sequence-to-function models to prioritize candidate regulatory variants for functional validation or clinical interpretation, and who need a concrete basis, available before any experiment is run, for deciding whether a prediction can inform a downstream translational or research decision or requires further evidence first. |

## 2. Methods

### 2.1. Study design and data sources

Training and internal-test data were ENCODE ATAC-seq on K562 (experiment ENCSR868FGK, GRCh38), using the fold-change-over-control bigWig track (ENCFF019IPA) and the replicated narrowPeak peak set (ENCFF057UYP, 258,895 peaks on primary chromosomes) [18]. The reference genome was UCSC hg38.2bit. Cross-assay validation data were lentiMPRA, an episomal massively parallel reporter assay [19], in the K562 arm of a joint K562/HepG2/WTC11 library (ENCODE experiment ENCSR203UFY) [20], using the per-element BED file with genomic coordinates (ENCFF802FUV, 53,989 elements before filtering). Perturbation-validation data were the PKLR-promoter arm of a 21-locus saturation-mutagenesis MPRA panel [21], the only locus in that panel assayed in K562 cells, sourced from the study’s OSF data repository (Supplementary Table S5).

### 2.2. Sequence-window construction

Positive windows were 2,048 bp regions centered on each ENCODE K562 ATAC-seq peak; matched negative windows of the same width were sampled from the same chromosomes with any peak overlap rejected. This produced 517,790 windows (258,895 positive, 258,895 negative). Chromosomes chr8 and chr9 were held out entirely as an internal test set (42,844 windows), never used for training, hyperparameter search, or reference-population calibration. Sequence was read on demand from hg38.2bit at train and inference time and one-hot encoded, with ambiguous or N bases encoded as an all-zero vector.

### 2.3. Model architecture and hyperparameter search

A convolutional architecture was chosen because chromatin accessibility at this scale is governed by local sequence composition and transcription-factor motifs occurring within a few hundred base pairs, the receptive-field structure a stacked convolutional kernel is built to learn directly. Convolutional networks form one of the standard architecture classes surveyed for sequence-to-function prediction in regulatory genomics, alongside transformer- and state-space-based architectures that extend the same task to longer-range genomic context; representative examples include the convolutional models Basset and ChromBPNet and the longer-range architectures Enformer, Borzoi, and Hyena/Mamba-based state-space models [7]. The model is a compact one-dimensional convolutional network taking a one-hot-encoded 2,048 bp window as input and predicting a single continuous accessibility value. Architecture and optimizer hyperparameters were selected by a 30-trial Optuna [22] search on one chromosome-grouped fold, searching learning rate, weight decay, batch size, convolutional channel profile, kernel width, and dropout, using a reduced epoch budget and a subsample of the training pool. The best configuration (channel profile 64-96-128, kernel width 9, dropout 0.0953, learning rate 0.001177, weight decay 4.42e-5, batch size 32) was used for the full-scale ensemble.

### 2.4. Ensemble training and cross-validation

A five-fold chromosome-grouped ensemble was trained over all windows not on the held-out chromosomes, with one model per fold and no separate retraining step. Label standardization (mean and standard deviation) was fit independently for each fold from that fold’s own training indices only, to avoid leaking the validation chromosome’s label distribution into that fold’s preprocessing. Each fold’s checkpoint stores its own architecture parameters and its own label-scaling statistics; at inference time, each ensemble member’s prediction is de-standardized with its own checkpoint’s statistics before averaging in raw units.

### 2.5. Internal-test evaluation

The deployed five-model ensemble was scored on the chr8/chr9 holdout. Ensemble mean and standard deviation per window were computed after per-model de-standardization; a training-pool reference sample (n = 5,000) was used to compute a population standard deviation for the consensus ratio described below. Performance was summarized by Spearman correlation, root-mean-square error, mean absolute error, bias, and skill relative to a constant-value null predictor, with a 95% confidence interval on skill obtained by bootstrap.

### 2.6. Occlusion attribution and the K562 motif shell

Attribution for one ensemble member per test window (the member with the highest validation Spearman correlation) was computed by systematic occlusion across 64 bins per 2,048 bp window (32 bp per bin), with mean absolute attribution per bin used as the localization signal. Occlusion-based attribution was used in preference to gradient-based methods, which carry documented artefacts for genomic sequence models unless explicitly corrected [23]. A curated panel of ten K562-relevant transcription-factor motifs (JASPAR position frequency matrices for GATA1, GATA1::TAL1, TAL1::TCF3, KLF1, NFE2, MAF::NFE2, GATA2, RUNX1, MYB, and STAT5A::STAT5B, representing the erythroid, megakaryocytic, and BCR-ABL-signalling transcriptional program K562 biology is organized around) was scanned against each window on both strands at a per-motif false-positive rate of 0.001. A window’s coherence score is the fraction of high-attribution occlusion bins that overlap at least one motif hit from this panel; windows with no resolvable motif shell were excluded from coherence-dependent analyses rather than assigned a default scenario. Attribution was computed from a single ensemble member rather than aggregated across all five because coherence here evaluates attribution against an external biological prior, not inter-model agreement; the consensus axis captures ensemble disagreement separately, and motif-based sequence representations of this kind carry substantial, independently demonstrated cell-type-specific regulatory information [24].

### 2.7. Motif-span causal occlusion and context-sensitivity analysis

To test whether the fixed 32-bp occlusion scheme obscured motif-specific effects, we performed a complementary motif-span perturbation analysis using the same trained checkpoint and K562 JASPAR motif panel. Contiguous motif-covered bases were treated as the perturbation unit and masked using the same replacement convention as the original occlusion analysis. For each sequence, the prediction change after masking the motif shell was compared with the mean absolute prediction change produced by five equal-length random control masks placed elsewhere in the same sequence. Causal coherence was defined as

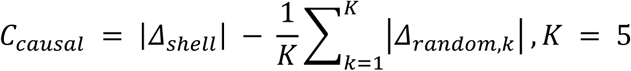

This matched control was used to distinguish motif-specific dependence from the generic effect of masking a larger amount of sequence. For sequences containing multiple motif instances, context dependence was evaluated by comparing the absolute prediction change from joint masking of nearby motif instances with the absolute summed effect of masking the same instances individually. Motif instances separated by no more than 24 bp were grouped into local modules; this distance was specified as an operational proximity threshold rather than a biologically derived interaction boundary. A module of n instances was classified as superadditive when

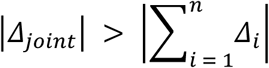

The causal-coherence analysis was performed on the held-out test set for reliability comparison and on the full 517,790-window dataset for design-facing mechanistic analysis. Reliability conclusions remained based on the held-out chromosomes. Throughout this subsection, “causal” denotes a controlled intervention on the model’s input sequence, not a claim of causal inference about transcription-factor biology.

### 2.8. Ensemble consensus and the four-scenario trust taxonomy

Ensemble consensus, disagreement among independently trained ensemble members as a practical estimate of predictive uncertainty [25], was defined as cv_ratio, the ratio of ensemble standard deviation to the population standard deviation from the training-pool reference sample. That reference sample is the same artificially balanced accessible/inaccessible mixture used for training (matched positive/negative sampling, see Sequence-window construction), so cv_ratio is calibrated against a balanced background population rather than genome-wide window frequencies; this is the appropriate choice for the training and internal-test analyses reported here, since all downstream evaluation in this study uses the same balanced construction. Consensus and coherence cutoffs were calibrated on an independent pool sample never used as the internal test set. Crossing binarized consensus and coherence at these cutoffs produces four scenarios: A (high consensus, high coherence), B (high consensus, low coherence), C (low consensus, high coherence), and D (low consensus, low coherence). Enrichment factor for a scenario at a given absolute-error threshold is that scenario’s precision (fraction of windows below the threshold) divided by the population base rate at the same threshold; a scenario is flagged unstable when its sample size falls below a minimum count. No scenario’s enrichment estimate was flagged unstable at the sample sizes reported here.

### 2.9. Applicability-domain distance

A pooled pre-head embedding (128-dimensional) was extracted for each window from the attribution checkpoint. Applicability-domain distance, a representation-space out-of-distribution measure of the kind reviewed for bioinformatics applications more broadly [26], is the cosine nearest-neighbor distance from a query window’s embedding to a 5,000-window training-pool reference sample. The applicability-domain cutoff was calibrated as the 95th percentile of the reference pool’s own leave-one-out self-distances. Quantile stratification bins the test set into percentile bands of this distance and reports mean absolute error and root-mean-square error per band.

### 2.10. Composition-divergence and dinucleotide-shuffle novelty axes

Composition divergence was defined as the Jensen-Shannon divergence between the 4-mer frequency distribution of each query window and that of the training-pool reference set. K-mer frequency profiles provide a simple sequence-level representation of genomic composition that is independent of the trained model [27]. The same 5,000-window reference pool used for applicability-domain estimation was used here. Dinucleotide-preserving shuffles were generated for 3,000 test windows using the Altschul-Erickson sequence-randomization procedure, which preserves dinucleotide frequencies while disrupting higher-order sequence organization [28]. The implementation follows the same general procedure used by sequence-shuffling utilities such as MEME Suite’s fasta-dinucleotide-shuffle and genomics workflows based on shuffled reference sequences. Candidate windows containing an N base were skipped and resampled. Applicability-domain distance, ensemble-predicted accessibility, and motif-shell coverage were then compared between each original sequence and its shuffled counterpart using paired Wilcoxon signed-rank tests.

A third candidate novelty definition tested whether repeat-derived sequence content identifies low-trust predictions. Per-window repeat-element overlap was computed from the UCSC hg38 RepeatMasker track, with overlapping repeat intervals merged per chromosome before computing overlap, and a window was classified repeat-derived if repeat-annotated bases exceeded 50% of its length. Per-window GC content was computed directly from the reference genome. Logistic regression modeled Scenario D and Scenario A membership as a function of repeat-derived status, applicability-domain distance (standardized to one standard deviation), and GC content, first on the chr8/chr9 held-out test set alone and then, as a stability check, on the full 517,790-window dataset (held-out test set and training pool combined).

### 2.11. Cross-assay lentiMPRA validation

lentiMPRA elements were obtained from the joint K562, HepG2, and WTC11 regulatory-element library [20], in which a common set of candidate regulatory elements was tested across all three cell types. Elements were filtered to primary chromosomes, retaining 53,950 of 53,989 sequences, and grouped according to their original design subgroup as K562-native, HepG2-designed, or WTC11-designed candidates. All elements analysed here were evaluated using the K562 lentiMPRA measurements [20]. Each element was mapped to a fixed-width sequence window centred on its midpoint. Occlusion attribution from the same checkpoint used for the coherence analysis was then reduced to a single element-level value by averaging the absolute attribution across bins overlapping the element. For the top 10% of elements by measured activity within each subgroup, we calculated the Spearman correlation between attribution and measured activity and a rank-based partial correlation controlling for the model’s predicted signal. We also fitted an ordinary least-squares model with measured activity as the response and standardized predicted signal and attribution as predictors, allowing the independent association of attribution with activity to be evaluated after accounting for predicted activity.

### 2.12. PKLR saturation-mutagenesis perturbation test

For each of 1,776 (24h) and 1,794 (48h) single-nucleotide variants at the PKLR promoter, a single fixed 2,048 bp reference window centered on the locus was scored once through the deployed five-model ensemble, using the accuracy-oriented, not attribution-oriented, ensemble prediction. Each variant’s predicted allele-substitution effect was computed as the ensemble’s predicted value for the alternate-allele sequence minus the reference-allele sequence on that same window. This predicted delta was compared against the measured log2 RNA-to-DNA ratio [21] by Spearman correlation, separately for each timepoint. The reference window’s own applicability-domain distance was computed once at the locus level, since a single-base substitution does not meaningfully move a 2,048 bp window’s embedding.

### 2.13. Statistical analysis

All correlations were Spearman rank correlations unless stated otherwise. Group comparisons used the Mann-Whitney U test for independent groups [29] or the paired Wilcoxon signed-rank test for paired observations [30], as appropriate. All tests were two-sided. Partial correlations controlling for a third variable were computed on ranks. Predictive skill relative to the constant null was defined as the difference between the null-model and ensemble mean absolute errors,

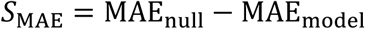

where the null predictor assigns every test window the mean accessibility of the training pool. Positive skill values therefore indicate lower error than the null predictor. A 95% confidence interval for the skill estimate was obtained by bootstrap resampling. No multiple-testing correction was applied across the distinct, pre-specified hypotheses reported in each results section. Thresholds and reference populations were fixed before each test was run. For the multivariable partial-correlation analyses, each variable was rank-residualized against the joint covariate set comprising target magnitude, applicability-domain distance, and motif-instance count before the residual correlation was calculated. Sensitivity analyses expanded this adjustment set to include GC content, repeat-derived content, and KLF1 instance density (Supplementary Table S13).

## 3. Results and Discussion

### 3.1. A convolutional ensemble learns transferable K562 chromatin-accessibility signal

A five-model chromosome-grouped ensemble was trained to predict K562 ATAC-seq accessibility from 2,048 bp DNA windows, with chr8 and chr9 reserved as an internal test set. Hyperparameter optimization produced a narrow validation Spearman range of 0.73–0.76 across all 30 trials, indicating that performance was not dependent on a single sharply optimal configuration. Cross-validated performance was consistent across folds, with a mean validation Spearman correlation of 0.7475 (SD 0.0154) (Supplementary Table S1, Figure 1B). On the held-out chr8/chr9 test set (n = 42,844), the ensemble reached Spearman *ρ* = 0.782, RMSE = 0.641, and MAE = 0.508 (Supplementary Table S6, Figure 1A). The MAE-based skill relative to the constant-value null predictor was 0.328 (95% CI [0.322, 0.333]). These results establish that the ensemble learned sequence-to-accessibility relationships that transferred to chromosomes excluded from training, providing the test bed for the prediction-specific reliability analyses that follow.

**Figure 1.**
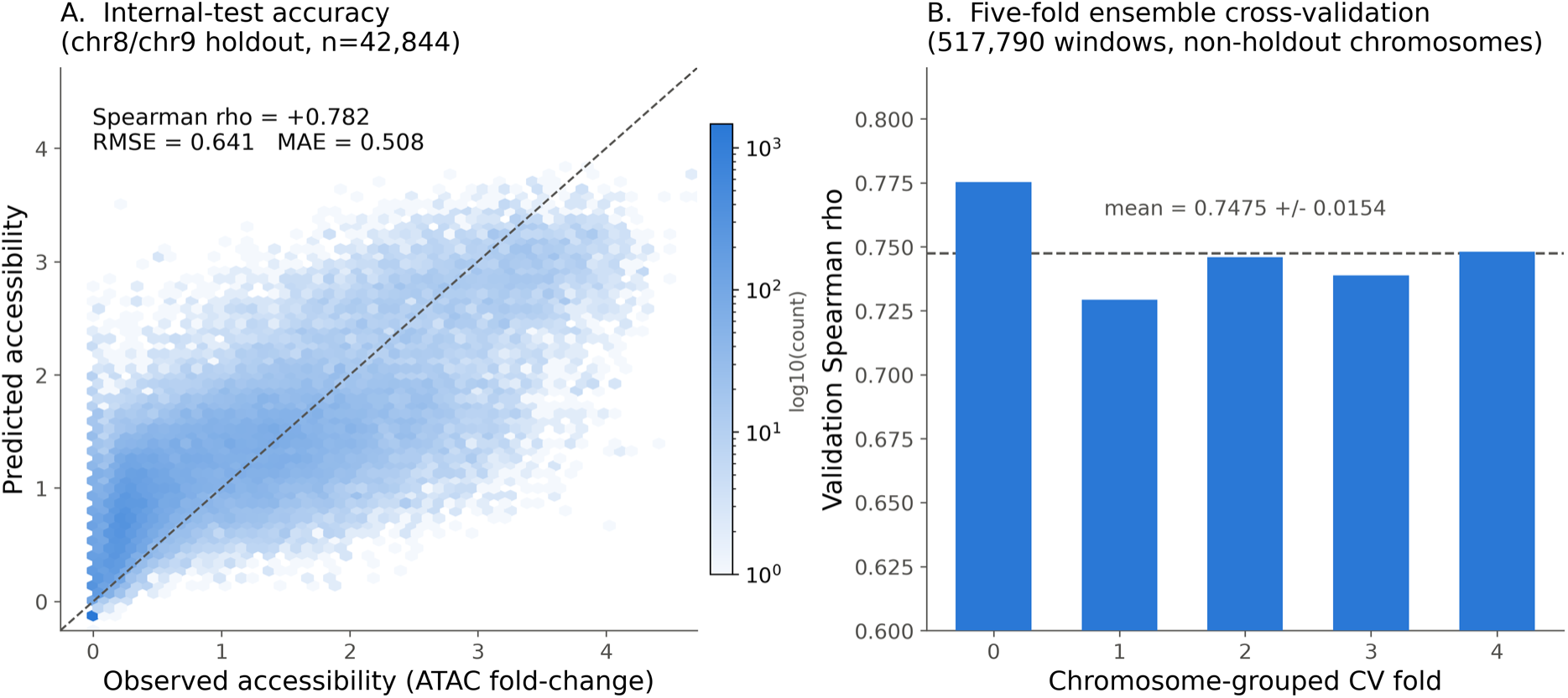
A five-model CNN ensemble learns transferable K562 chromatin-accessibility signal. (A) Predicted versus observed accessibility on the chr8/chr9 internal-test holdout (n=42,844), hexbin density. (B) Per-fold validation Spearman correlation across the five-fold chromosome-grouped ensemble.

### 3.2. Ensemble consensus and attribution coherence jointly separate prediction reliability

Each held-out test window was assigned to one of four scenarios by crossing two binary inference-time signals. Ensemble consensus captured agreement among the five ensemble members relative to a calibrated cutoff, while attribution coherence measured whether occlusion-based attribution overlapped a curated panel of ten K562-relevant transcription-factor motifs scanned on both strands. The panel comprised GATA1, GATA1::TAL1, TAL1::TCF3, KLF1, NFE2, MAF::NFE2, GATA2, RUNX1, MYB, and STAT5A::STAT5B. Scenario A (high consensus, high coherence) covered 10.06% of the 41,379 test windows that resolved a motif shell; Scenario B (high consensus, low coherence) covered 22.99%; Scenario C (low consensus, high coherence) covered 17.76%; and Scenario D (low consensus, low coherence) covered 49.20% (Table 1, Figure 2A).

**Figure 2.**
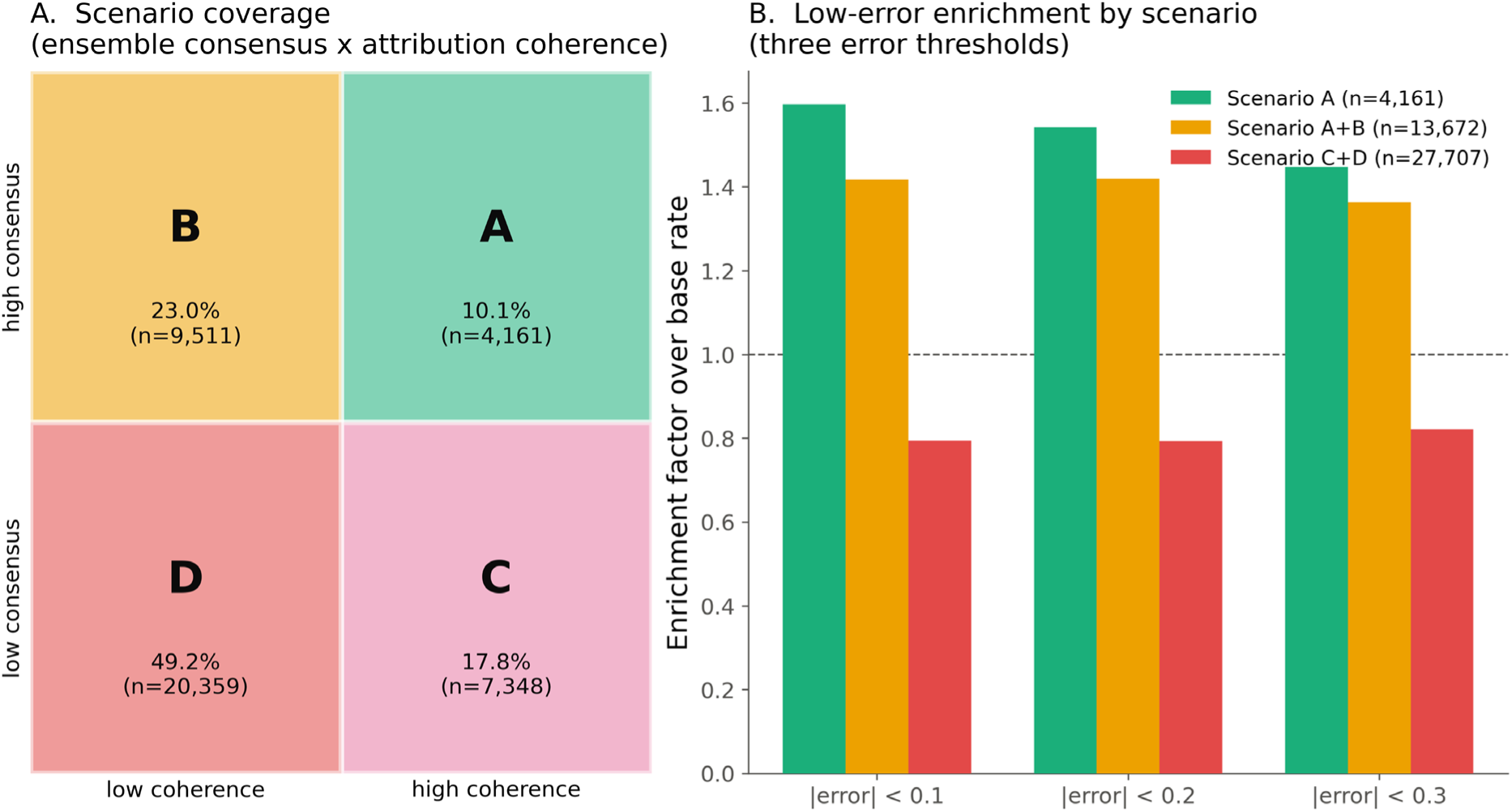
Ensemble consensus and attribution coherence jointly separate prediction reliability. (A) Scenario coverage from crossing binarized consensus and coherence. (B) Low-error enrichment factor for Scenario A, A+B, and C+D at three error thresholds.

**Table 1.**
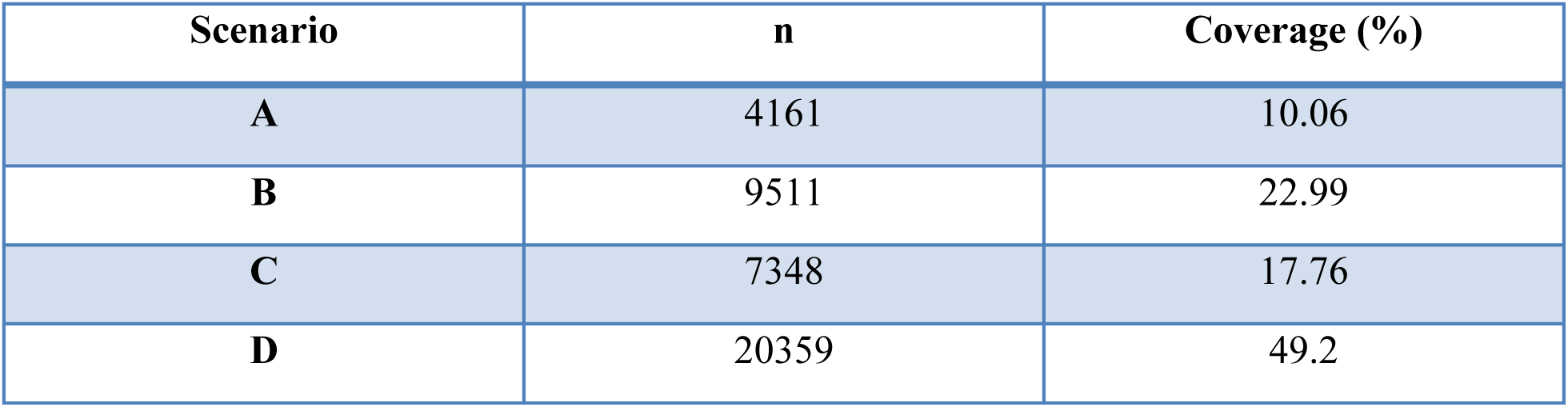
Trust-taxonomy scenario coverage on the internal test set.

| Scenario | n | Coverage (%) |
| --- | --- | --- |
| **A** | 4161 | 10.06 |
| **B** | 9511 | 22.99 |
| **C** | 7348 | 17.76 |
| **D** | 20359 | 49.2 |

A parallel full-dataset run of the same bin-overlap taxonomy (517,790 windows, 502,898 with a resolvable motif shell) produced a closely similar scenario distribution to the held-out chromosomes (Supplementary Table S10), supporting the stability of the qualitative trust taxonomy across the analysed sequence collection.

Low-consensus predictions accounted for most evaluable windows, with Scenario D alone comprising 49.2% of the test set. Figure 2 shows how this scenario structure translates into enrichment for low-error predictions.

Scenario A and the combined Scenario A+B population were enriched for low-error predictions relative to the combined Scenario C+D population at every error threshold tested. At the ∣error∣<0.1 threshold, the enrichment factor over the population base rate was 1.596 for Scenario A, 1.417 for A+B, and 0.794 for C+D; the same ordering held at the 0.2 and 0.3 thresholds (Figure 2B). The enrichment pattern in Figure 2 shows that high-consensus predictions are generally more reliable than low-consensus predictions, but it does not isolate the additional contribution of attribution coherence. Restricting the analysis to Scenarios A and B holds consensus status constant and therefore tests whether coherence adds further discrimination within the high-consensus population. Coherence further separated error within the high-consensus population (mean absolute error 0.396 in Scenario A versus 0.435 in Scenario B, Mann-Whitney p = 9.6e-10), with Scenario A’s precision exceeding Scenario B’s by 1.09-to 1.19-fold across the 0.1-0.3 error thresholds (Table 2, Figure 3).

**Figure 3.**
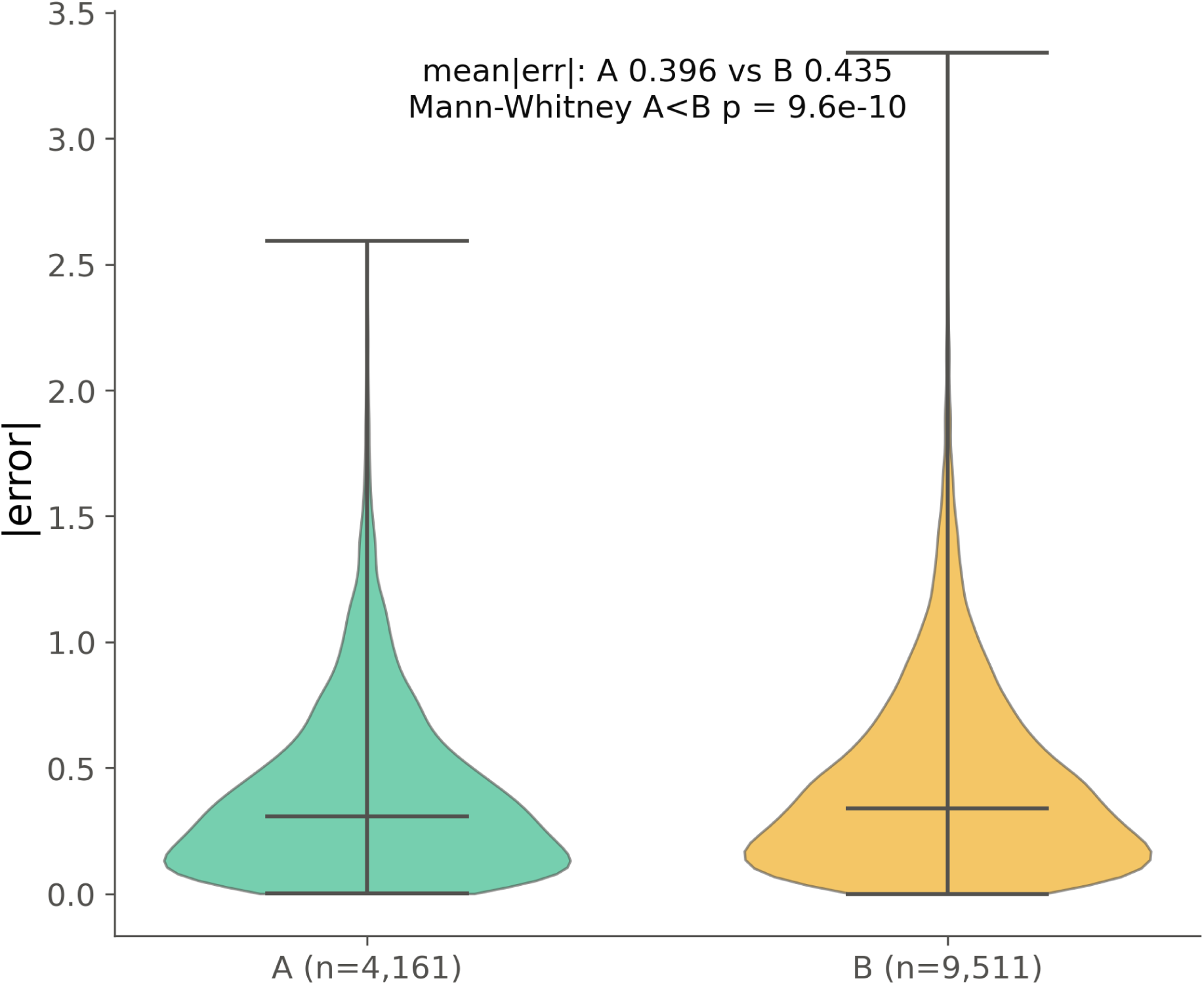
Attribution coherence separates prediction error within the high-consensus population.

**Table 2.**
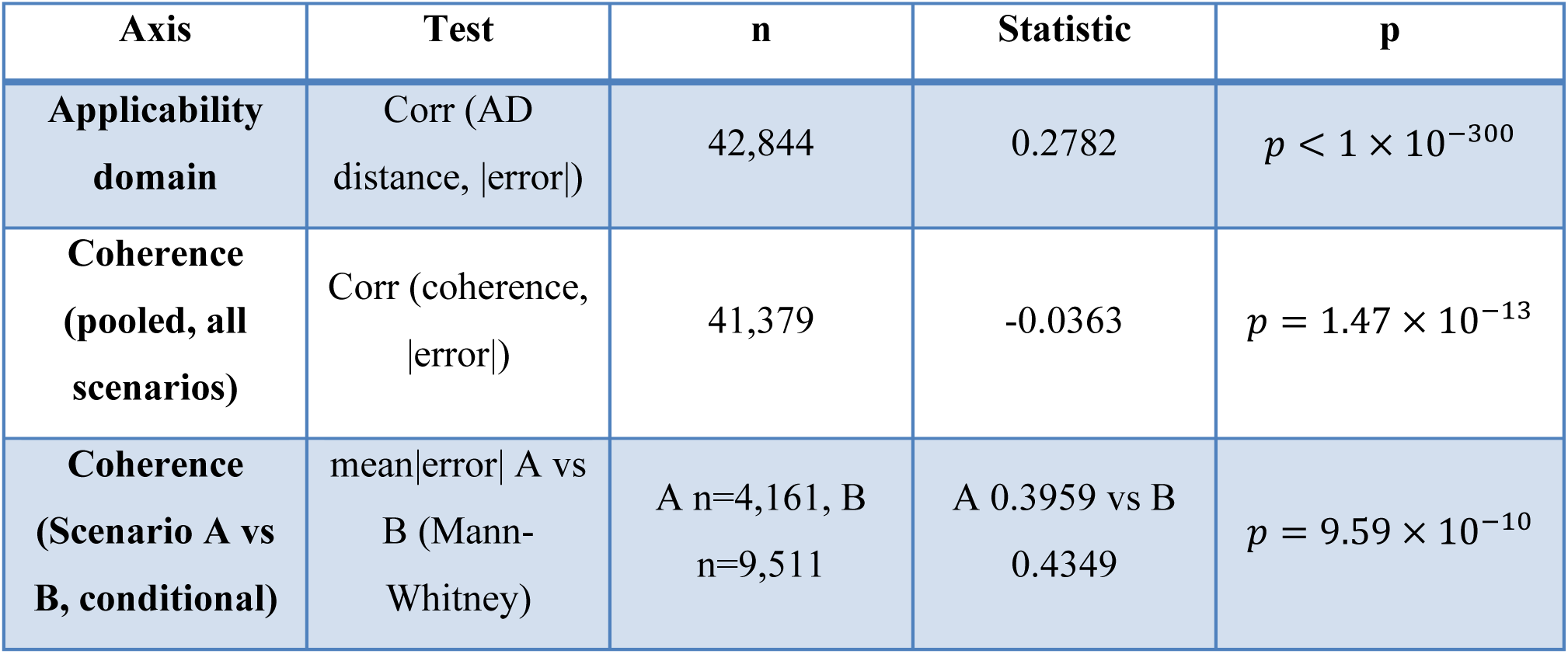
Validation of the applicability-domain and coherence axes against held-out error.

### 3.3. Applicability-domain distance maps a competence boundary that predicts increasing error

Applicability-domain distance was computed as the cosine nearest-neighbor distance between each test-window embedding and a 5,000-window reference sample drawn from the training pool. Across the held-out test set, greater applicability-domain distance was associated with larger absolute prediction error (Table 2), the strongest single-axis relationship among the three inference-time signals evaluated (Table 2, Figure 4A). Stratifying the test set into six applicability-domain percentile bands produced a monotonic error gradient. Mean absolute error increased from 0.355 in the nearest 20% of windows to 0.683 in the farthest 5% (Supplementary Table S2, Figure 4B). The calibrated 95th-percentile cutoff classified 4.9% of the test set as out-of-domain. MAE was 0.500 for in-domain windows and 0.681 for out-of-domain windows (Figure 4C).

**Figure 4.**
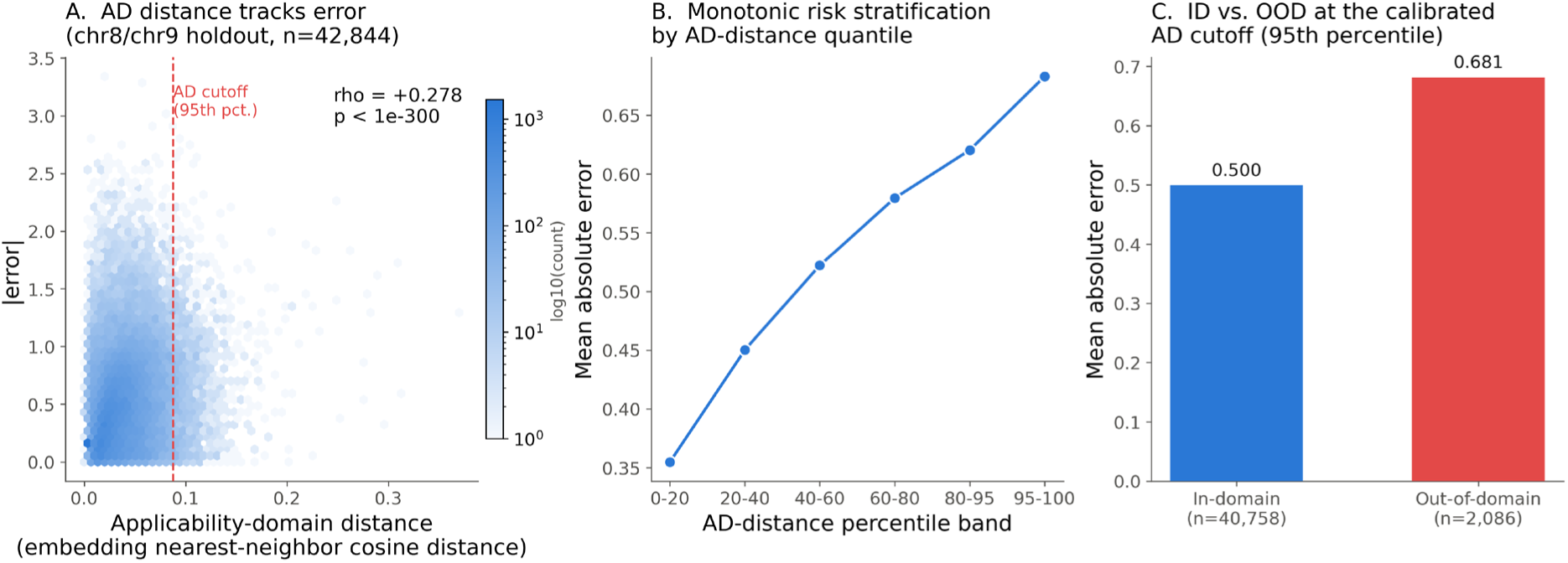
Applicability-domain (AD) distance maps a competence boundary that predicts increasing error. (A) AD distance versus |error| across the internal-test holdout. (B) Mean absolute error by AD-distance percentile band. (C) Mean absolute error, in-domain versus out-of-domain, at the calibrated 95th-percentile cutoff.

The contrast between the pooled and conditional coherence results also clarifies the relative roles of the trust signals. Coherence has little association with error when considered across all scenarios, whereas applicability-domain distance shows a substantially stronger global relationship with error. Figure 4 examines this applicability-domain relationship directly, showing whether the continuous association translates into an ordered increase in prediction error as sequences move farther from the training representation.

### 3.4. Learned applicability-domain distance captures sequence novelty

Unexpectedly, 4-mer composition divergence from the training pool was negatively associated with prediction error ( *ρ* = −0.180, *p* < 1 × 10^−300^, n = 42,844; Supplementary Table S4, Supplementary Figure S1B) and anti-correlated with embedding-space applicability-domain distance on the same test set ( *ρ* = −0.604, *p* < 1 × 10^−300^; Supplementary Figure S1A). Windows with more unusual k-mer composition therefore tended to have lower error, and sequences scored as compositionally unusual were often treated as relatively familiar by the learned embedding. Dinucleotide-preserving shuffling provided a complementary test. Shuffling 3,000 test windows while preserving dinucleotide composition increased applicability-domain distance in 80.5% of windows (Wilcoxon *p* = 1.8 × 10^−280^), decreased predicted accessibility (Wilcoxon *p* = 8.1 × 10^−20^), and decreased motif-shell coverage (Wilcoxon *p* = 4.4 × 10^−14^) (Supplementary Figure S1D-F). These results indicate that preserving coarse sequence composition does not preserve the model’s learned representation of regulatory structure.

The divergence between composition-based and embedding-based novelty suggests that applicability-domain distance captures more than unusual base composition. Regulatory activity depends on features such as motif spacing, orientation, and combinatorial arrangement, which can be disrupted while overall sequence composition is retained [31]. The present analysis tests one composition-based alternative, so other sequence-level novelty measures may still prove informative. Repeat-derived sequence content provided a further test of whether an intuitive biological descriptor identifies low-trust predictions. On the chr8/chr9 holdout, windows with more than 50% repeat-derived sequence had lower odds of Scenario D after adjustment for applicability-domain distance and GC content (OR = 0.78, 95% CI 0.747-0.812) and higher odds of Scenario A (OR = 1.39, 95% CI 1.298-1.488; n = 41,375). The full 517,790-window analysis produced nearly identical odds ratios of 0.79 and 1.39, respectively (Supplementary Table S9), indicating that the result was not explained by repeat-heavy windows having participated in model fitting. Repeat content therefore did not identify a low-trust region of this model at the >50%-of-window threshold tested. A plausible explanation is that common repeat classes recur extensively across the genome and are therefore well represented during training, although the present analysis does not directly test that mechanism.

### 3.5. Independent assay validation tests explanation generalizability

Attribution coherence separated prediction error within the high-consensus population, but this does not establish whether the same attribution signal tracks regulatory activity measured by an independent assay. We tested this using K562 lentiMPRA data for the top 10% of elements by measured activity in three design subgroups comprising K562-native elements (n = 1,788), HepG2-designed elements (n = 1,806), and WTC11-designed elements (n = 1,802). Attribution was compared with measured activity using partial Spearman correlation while controlling for the ensemble’s predicted accessibility. The relationship was strongly subgroup-dependent (Table S3, Figure S2). Partial correlation was negligible for K562-native elements (*ρ*_partial_ = 0.020, p = 0.257), weakly negative for HepG2-designed elements (*ρ*_partial_ = −0.028, p = 0.0012), and positive for WTC11-designed elements ( *ρ*_partial_ = 0.193, *p* = 3.3 × 10^−12^). Attribution coherence therefore did not transfer uniformly from the chromatin-accessibility task to activity measured by lentiMPRA.

This result distinguishes explanation usefulness within the model’s original prediction setting from explanation validity across experimental contexts. lentiMPRA measures activity in an episomal reporter context, whereas the model was trained on chromosomal accessibility, and these assay contexts can differ systematically [32]. A similar distinction has been reported in molecular taste prediction, where dataset-specific structural associations can appear mechanistically meaningful despite reflecting underlying composition biases [14, 33]. In RegTrust-XAI, the subgroup-dependent lentiMPRA result provides the corresponding empirical caution in regulatory genomics. Attribution coherence can help identify more reliable predictions within the model’s native setting, but biological plausibility alone should not be interpreted as evidence that the attributed feature transfers mechanistically across assays.

### 3.6. PKLR saturation-mutagenesis validates variant-effect predictions

Single-nucleotide saturation-mutagenesis MPRA data at the PKLR promoter in K562 cells [21], the only locus in the 21-element panel assayed in the same cell type used here, provided a direct perturbation test. For each of 1,776 variants at 24 h and 1,794 variants at 48 h, the predicted allele-substitution effect was calculated as the ensemble prediction for the alternate allele minus that for the reference allele within the same 2,048 bp window. Predicted allele-substitution effects correlated with experimentally measured effects at both 24 h ( *ρ* = 0.227, *p* = 4.1 × 10^−22^) and 48 h (*ρ* = 0.235, *p* = 6.1 × 10^−24^) (Table S4, Figure 5). The reference window had an applicability-domain distance of 0.0068, well below the calibrated cutoff of 0.0875, placing the locus within the model’s learned applicability domain.

**Figure 5.**
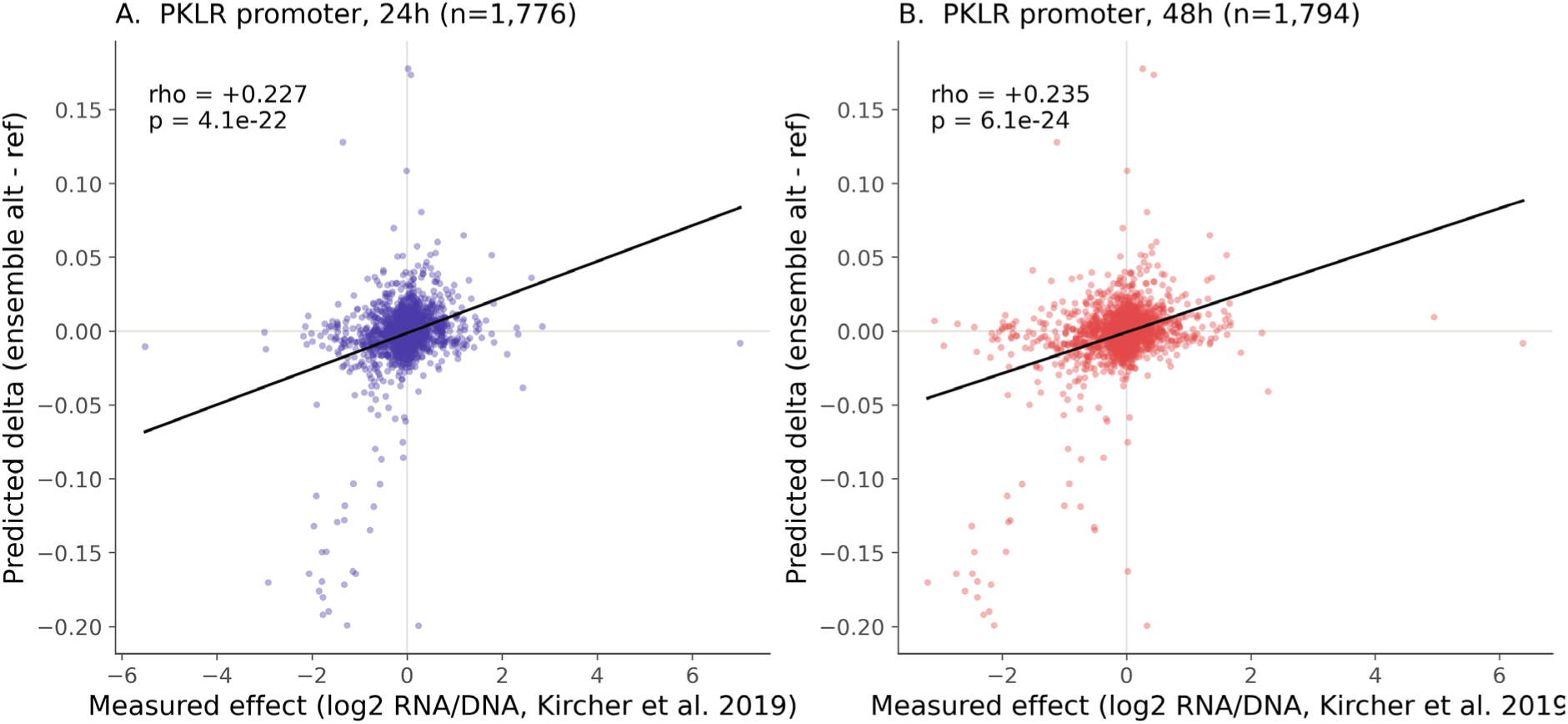
Predicted single-nucleotide effects agree with saturation-mutagenesis measurements at the PKLR promoter [21]. Predicted allele-substitution effect (ensemble alt minus ref) versus measured effect (log2 RNA/DNA), at (A) 24 hours and (B) 48 hours post-transfection.

The PKLR analysis shows that the model captures a reproducible component of measured single-nucleotide regulatory effects at this locus. Unlike the lentiMPRA comparison, this analysis evaluates predictions against direct sequence perturbations rather than cross-assay associations. The result therefore provides perturbation-level support for variant-effect prediction at this locus, while genome-wide generalization remains to be established across additional regulatory elements and cell types. Additional perturbation datasets will be required to determine how broadly this relationship generalizes. Large-scale variant-effect models such as AlphaGenome aim to predict regulatory consequences across the genome, whereas RegTrust-XAI addresses whether an individual prediction lies within an empirically supported competence region before experimental confirmation is available.

### 3.7. Motif-specific perturbation reveals context-dependent regulatory attribution

The fixed-bin coherence metric is operationally convenient but does not align exactly with motif boundaries: a 32-bp bin can contain both a motif and unrelated flanking sequence, and a motif can span more than one bin. We therefore repeated the explanation analysis using motif-span-exact causal occlusion with equal-length random masking controls.

On the held-out high-consensus population, causal coherence showed a stronger unadjusted association with prediction error than bin-overlap coherence. However, this apparent advantage was substantially attenuated after accounting for target magnitude, applicability-domain distance, and the number of motif instances. The same pattern was reproduced at full-dataset scale. Across the full high-consensus population (n = 164,585), the correlation between causal coherence and absolute error changed from *ρ* = −0.1702 to partial *ρ* = −0.0423 after adjustment, whereas bin-overlap coherence changed from *ρ* = −0.0476 to partial *ρ* = −0.0277. Thus, the biologically more targeted perturbation did not provide a clearly superior independent error-discrimination signal and was not adopted as a replacement for the operational coherence axis.

The same analysis nevertheless revealed a strong model-level non-additive pattern. Across 4,703,837 multi-instance motif modules in the full dataset, 90.4% showed a superadditive joint-knockout effect, closely reproducing the 91.1% rate observed in the held-out analysis. This indicates that the model’s response to a motif depends strongly on neighbouring regulatory sequence context, so an attribution assigned to an isolated motif should not be interpreted as an intrinsic, context-independent property of that motif.

This pattern of context-dependent motif contribution is consistent with prior sequence-to-function studies showing that transcription-factor motif effects can depend on neighbouring motifs, spacing, orientation, and flanking sequence [5, 34]. Related perturbation-based interpretation approaches have likewise shown that genomic deep-learning models can encode dependencies among sequence features rather than acting on isolated motifs alone [35, 36]. Regulatory grammar is not universally strongly cooperative, however: a large-scale reporter-assay study found that transcription factors generally act additively with comparatively weak grammar in most tested contexts [37]. The present result is therefore interpreted as a property of the regulatory dependencies learned for this K562 accessibility model, not evidence for a universal rule of cis-regulatory grammar.

### 3.8. Implications for trustworthy regulatory sequence modelling

The three inference-time axes captured different and only partially overlapping aspects of prediction reliability. Consensus and applicability-domain distance produced the strongest separation in prediction error, while attribution coherence added a smaller but complementary signal within the high-consensus population. High consensus indicates stability across independently trained models, low applicability-domain distance indicates that a sequence lies within a familiar region of learned representation space, and high coherence indicates that the attribution overlaps regulatory features relevant to K562 biology. Predictions supported by all three signals therefore have the strongest inference-time evidence, whereas predictions outside the applicability domain should be treated as extrapolative and low-consensus predictions warrant greater caution. Attribution coherence remains distinct from causal validation because motif-grounded attribution does not establish that the highlighted feature is mechanistically responsible for the observed regulatory effect, as demonstrated by the lentiMPRA and PKLR analyses.

The sequence-novelty analyses further clarify what model familiarity represents. Neither unusual 4-mer composition nor high repeat-derived sequence content identified the low-trust population, whereas dinucleotide shuffling, which disrupts higher-order sequence organization while preserving composition, moved sequences farther from the learned applicability domain. Model familiarity therefore appears to depend more strongly on sequence structure captured in the learned embedding than on biological descriptors assumed in advance to indicate novelty. This distinction also separates data familiarity from representational sufficiency. Repeated sequence patterns may be extensively represented during training, whereas information absent from the model input cannot be recovered simply through greater familiarity with the represented features. Applicability-domain and consensus estimates address the former problem, while the latter requires reconsideration of the model representation itself. These findings address several limitations recently identified in regulatory-genomics modelling. Graded distribution-shift evaluation, calibrated uncertainty, and perturbational validation have been identified as unresolved requirements for developing more reliable sequence-to-function models [7]. Related studies have shown that genomic models can lose accuracy in cell-type-specific accessible regions [9] and that sequence-to-function performance can collapse for unseen genes despite strong performance elsewhere [10].Related challenges also arise in population-genomic inference, where recent methods such as TRACE infer archaic ancestry from reconstructed ancestral recombination graphs and show that reliability can depend strongly on the quality of the inferred genealogy and demographic setting, despite high precision under validated conditions [38]. RegTrust-XAI addresses the complementary problem of identifying elevated prediction risk before experimental ground truth becomes available.

The relationship between explanation coherence and prediction error was also conditional on applicability-domain status. Outside the learned applicability domain, both coherence formulations showed little additional pooled error discrimination. This does not mean coherence becomes meaningless outside the domain: OOD windows were already enriched for low-consensus Scenarios C and D, leaving a smaller, compositionally different high-consensus population for the A-versus-B comparison. The practical implication is hierarchical: applicability-domain status first determines whether the sequence lies within a region supported by the learned representation, after which consensus and motif-grounded coherence provide additional evidence about prediction stability and explanation grounding. Under the bin-overlap definition, OOD windows comprised approximately 4.9% Scenario A, 12.9% B, 21.1% C, and 61.1% D, compared with 10.2%, 23.3%, 18.1%, and 48.3%, respectively, within the applicability domain.

More broadly, the framework connects sequence-to-function benchmarking, ensemble-based uncertainty estimation, genomic model explainability, and experimental validation, which are often evaluated separately. Its contribution is to place prediction stability, explanation coherence, and applicability-domain distance within one framework as complementary but non-interchangeable forms of evidence for an individual prediction. The present analyses further show that these signals are not fully independent: motif-grounded coherence is partly coupled to applicability-domain position, and its additional error discrimination is strongly reduced among sequences already flagged as outside the learned domain. This is consistent with broader trustworthy-AI work showing that explainability alone does not establish trustworthiness and should be evaluated alongside complementary evidence [39]. A regulatory prediction should therefore be judged using multiple forms of evidence rather than predictive accuracy or biological plausibility alone.

The causal-occlusion analysis also shows why explanation granularity alone should not be equated with explanation validity. Masking motif boundaries more precisely raised the raw association between coherence and error, but most of that gain reflected shared variation with applicability-domain distance, target magnitude, and motif density. A more precise attribution unit therefore did not automatically yield a better trust signal.

## 4. Conclusion

Ensemble consensus and applicability-domain distance captured most of the error discrimination in the held-out test set, while motif-grounded coherence provided a smaller complementary signal among high-consensus predictions. A motif-span-specific perturbation analysis increased the unadjusted association between coherence and prediction error, but most of that gain was attenuated after accounting for applicability-domain distance, target magnitude, and motif density, so the causal formulation was not adopted as a replacement for the operational bin-overlap coherence measure. The same perturbation analysis revealed a separate mechanistic result, 90.4% of 4,703,837 multi-instance motif modules showed superadditive joint-knockout effects, indicating that regulatory attributions are strongly context-dependent rather than fixed properties of isolated motif instances. Explanation coherence also provided little additional error discrimination among sequences outside the learned applicability domain, a population already enriched for low-consensus predictions. Together, these results support a hierarchical interpretation of model trust, in which domain familiarity, ensemble stability, and biological grounding provide complementary, partially dependent evidence for prediction and explanation reliability. Several boundaries define how far these results generalize. The analyses are restricted to K562 chromatin accessibility, and transfer to other cell types and regulatory readouts remains to be established. Mutation-level validation was limited to 1,776–1,794 variants at a single PKLR locus. Applicability-domain distance is specific to the learned representation of the model, while motif-grounded coherence depends on the transcription-factor panel selected for the biological context. Both therefore require recalibration when the model architecture, cell type, or prediction task changes. For less-characterized cell types, the motif shell could be constructed dynamically from cell-type-specific accessibility, transcription-factor expression, ChIP-seq, or motif-enrichment data rather than from a fixed manually curated panel. The robustness of the present calibration was supported by sensitivity analyses. Varying the applicability-domain reference pool from 2,500 to 10,000 windows changed the distance-error correlation by less than 0.002, while alternative consensus and coherence thresholds preserved enrichment of Scenario A and A+B and depletion of C+D for low-error predictions (Supplementary Tables S7-S8).

Future evaluation should extend the framework to additional perturbation loci, cell types, and regulatory readouts, and should test consensus calibration against genome-representative rather than balanced reference populations. RegTrust-XAI is intended to support experimental prioritization by identifying predictions that are stable, biologically coherent, and within a model’s learned competence region before wet-lab resources are committed. It does not replace experimental validation, but provides a principled basis for deciding where that validation is most needed.

## Data and code availability

Codebase is available at https://github.com/MujeebOnawole/RegTrust-XAI. Raw data are publicly available from ENCODE (ENCSR868FGK, ENCSR203UFY), UCSC (hg38.2bit, RepeatMasker track), and an OSF repository for the saturation-mutagenesis dataset (Kircher et al., 2019; https://doi.org/10.17605/OSF.IO/75B2M).

## Acknowledgement

This work was supported by resources provided by The University of Queensland Research Computing Centre’s Bunya supercomputer.

