## Supplementary Information for "Trust-Aware Sequence-to-Function Modelling in Regulatory Genomics"

### Supplementary Figures

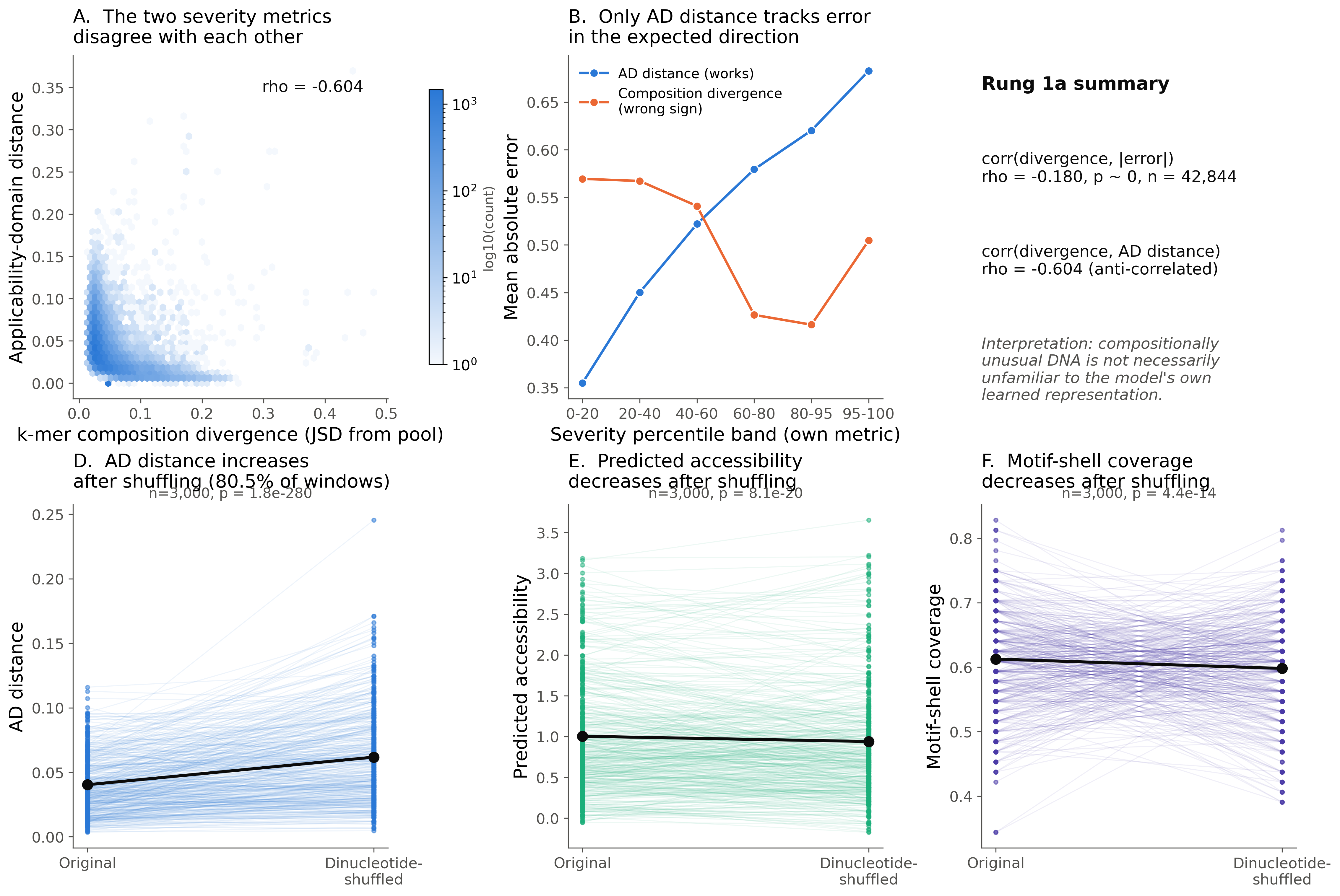

**Figure S1.** Composition-based novelty fails as a severity metric, while learned representation distance detects motif rearrangement. (A) k-mer composition divergence versus applicability-domain (AD) distance, same test windows (n=42,844). (B) Mean absolute error by severity percentile band for both candidate metrics. (C) Rung 1a summary statistics. (D-F) Paired original-versus-dinucleotide-shuffled comparison of AD distance, predicted accessibility, and motif-shell coverage (n=3,000). Referenced from the main text's "Learned applicability-domain distance captures sequence novelty beyond raw composition" section.

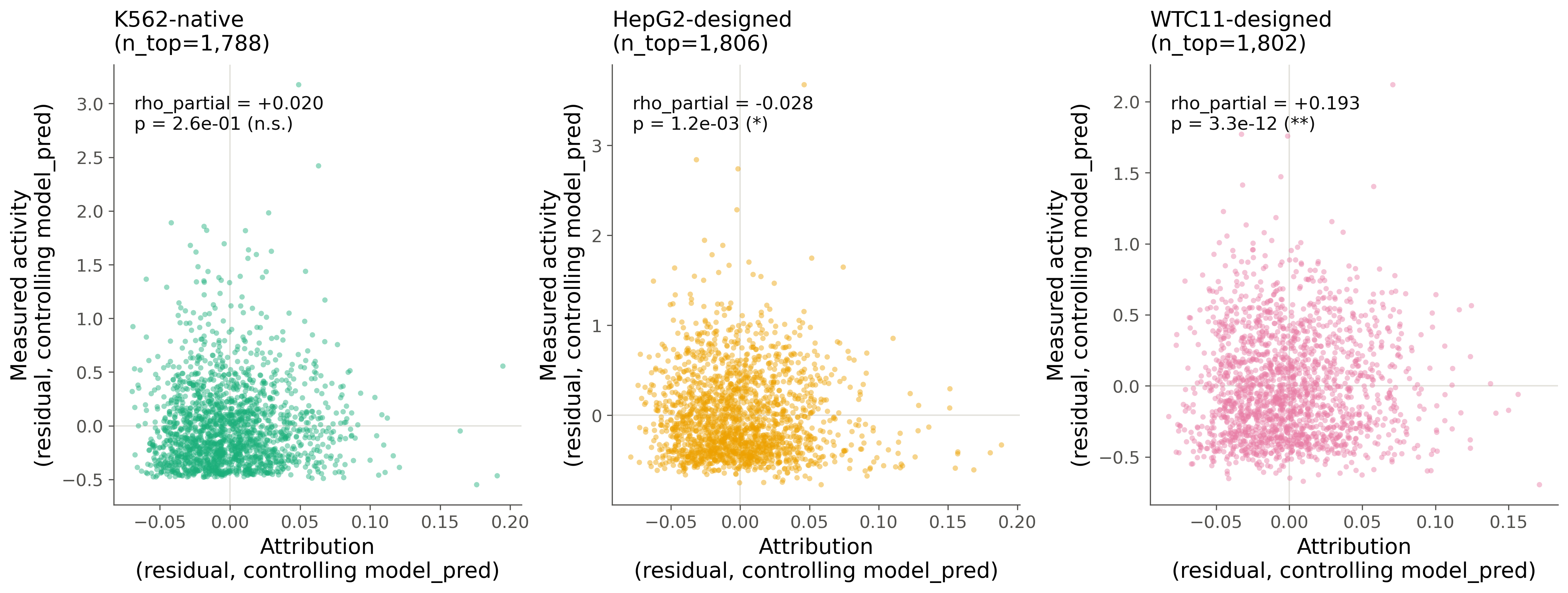

**Figure S2.** Explanation validity across an independent assay (lentiMPRA) is subgroup-dependent. Attribution and measured activity, both residualized on the ensemble's own predicted signal, for the top 10% of elements by activity in each of three subgroups: K562-native, HepG2-designed, and WTC11-designed. Referenced from the main text's "Attribution transfer to an independent assay is subgroup-dependent" section.

### Supplementary Tables

**Table S1.** Five-fold ensemble training, per-fold validation Spearman correlation. Referenced from the main text's "A convolutional ensemble learns transferable K562 chromatin-accessibility signal" section.

| Fold | Validation Spearman $\boldsymbol{\rho}$ | n train | n validation |
| --- | --- | --- | --- |
| 0 | 0.7753 | 376165 | 98781 |
| 1 | 0.7293 | 366127 | 108819 |
| 2 | 0.7459 | 393811 | 81135 |
| 3 | 0.7387 | 376525 | 98421 |
| 4 | 0.7481 | 387156 | 87790 |
| Mean +/- SD | 0.7475 +/- 0.0154 | - | - |

**Table S2.** Mean absolute error by applicability-domain distance percentile band, the tabular form of Figure 4B in the main text. Referenced from the main text's "Applicability-domain distance maps a competence boundary" section.

| AD-distance percentile band | n | MAE | RMSE |
| --- | --- | --- | --- |
| 0-20 | 8569 | 0.3549 | 0.4939 |
| 20-40 | 8569 | 0.4501 | 0.5693 |
| 40-60 | 8568 | 0.522 | 0.6362 |
| 60-80 | 8569 | 0.5794 | 0.6994 |
| 80-95 | 6426 | 0.6203 | 0.7508 |
| 95-100 | 2143 | 0.6829 | 0.8281 |

**Table S3.** Cross-assay attribution-activity correlation, lentiMPRA subgroups (raw and partial correlation, and the OLS attribution-coefficient decomposition). Referenced from the main text's "Attribution transfer to an independent assay is subgroup-dependent" section.

| Subgroup | n (top 10% by activity) | $\boldsymbol{\rho}$ (raw) | $\boldsymbol{\rho}$ (partial, controlling model_pred) | OLS attribution coefficient | OLS attribution p |
| --- | --- | --- | --- | --- | --- |
| K562-native | 1788 | -0.0314 | 0.0203 | -0.0306 | 0.257 |
| HepG2-designed | 1806 | -0.2481 | -0.0277 | -0.1012 | 0.00124 |
| WTC11-designed | 1802 | 0.0801 | 0.1932 | 0.2625 | 3.28e-12 |

**Table S4.** Summary of the graded distribution-shift and perturbation evaluation: rung 0 (chromosome holdout), rung 1 (sequence-novelty challenges, comprising three complementary diagnostics -- composition divergence (1a), dinucleotide shuffle (1b), and repeat-derived content (1c), the last reported in full, including all regression terms, in Table S9), rungs 2-3 (cross-assay lentiMPRA), and rung 4 (PKLR perturbation). Includes rungs already shown in the main text's Figures 2, 4, and 5, provided here as a single consolidated reference table.

| Rung | Definition | n | Metric | Value | p |
| --- | --- | --- | --- | --- | --- |
| 0. Chromosome holdout | chr8/chr9, same assay | 42844 | Spearman $\rho$ | 0.782 | - |
| 1a. Sequence-novelty: composition divergence | chr8/chr9, k-mer JSD from pool | 42844 | corr(divergence, \|error\|) | -0.18 | 1.01e-307 |
| 1b. Sequence-novelty: dinucleotide shuffle (AD check) | dinucleotide-shuffled windows | 3000 | Wilcoxon, AD distance increase | 0.805 | 1.85e-280 |
| 1c. Sequence-novelty: repeat-derived content (AD check) | chr8/chr9, >50% repeat-derived vs. AD-distance/GC-adjusted | 41375 | Logistic regression OR, Scenario D / Scenario A (full model in Table S9) | OR 0.78 (D), 1.39 (A) | see Table S9 |
| 2-3. Cross-assay MPRA | K562-native / HepG2 / WTC11-designed | 53950 | $\rho$ (partial), see Table 7 | -0.028 to +0.193 | - |
| 4. PKLR perturbation (24h) | Saturation mutagenesis, K562 | 1776 | corr(predicted delta, measured) | 0.227 | 4.07e-22 |
| 4. PKLR perturbation (48h) | Saturation mutagenesis, K562 | 1794 | corr(predicted delta, measured) | 0.235 | 6.12e-24 |

**Table S5.** Data sources used in this study. Referenced from the main text’s "Study design and data sources" section.

| Role | Source | Scale | Assembly |
| --- | --- | --- | --- |
| Training / internal test | ENCODE ATAC-seq, K562 (ENCSR868FGK) | 517,790 windows (258,895 positive + 258,895 matched negative); chr8/chr9 held out as internal test (n=42,844) | GRCh38 |
| Cross-assay validation | lentiMPRA, K562 arm (ENCSR203UFY; Agarwal et al. 2025) | 53,950 elements after contig filtering (k562_native 17,878 / hepg2_designed 18,057 / wtc11_designed 18,015) | GRCh38 |
| Perturbation validation | Saturation-mutagenesis MPRA, PKLR promoter, K562 (Kircher et al. 2019) | 1,776 variants (24h) / 1,794 variants (48h), single 469 bp locus | GRCh38 |

**Table S6.** Internal-test (chromosome holdout) performance of the deployed ensemble, the tabular form of Figure 1. Referenced from the main text’s "A convolutional ensemble learns transferable K562 chromatin-accessibility signal" section.

| Metric | Value |
| --- | --- |
| n | 42844 |
| Spearman $\boldsymbol{\rho}$ | 0.7817 |
| RMSE | 0.6413 |
| MAE | 0.5085 |
| Bias | 0.1447 |
| Skill vs. constant-null predictor | 0.3277 |
| Skill 95% CI | [0.3223, 0.3330] |
| Constant-null RMSE | 1.1582 |

**Table S7.** Sensitivity of the applicability-domain distance-error correlation and calibrated cutoff to the reference-pool size (2,500/5,000/10,000 windows; 5,000 is the value used throughout the main text). Referenced from the main text’s “Conclusion” section.

| Reference-pool size | Spearman $\boldsymbol{\rho}$ (AD distance vs \|error\|) | AD cutoff (95th percentile) | Out-of-distribution rate at cutoff (%) |
| --- | --- | --- | --- |
| 2,500 | 0.2772 | 0.0952 | 5.36 |
| 5,000 (baseline) | 0.2782 | 0.0875 | 4.87 |
| 10,000 | 0.2799 | 0.0813 | 4.68 |

**Table S8.** Sensitivity of Scenario A/A+B/C+D coverage and enrichment factor at the 0.1 absolute-error threshold to alternative consensus and coherence calibration percentiles. Baseline uses the 30th (consensus) / 70th (coherence) calibration percentiles described in Methods (Section 2.7); Consensus 20th/40th pct vary the consensus threshold with coherence fixed at the 70th percentile, and Coherence 60th/80th pct vary the coherence threshold with consensus fixed at the 30th percentile. Referenced from the main text’s “Conclusion” section.

| Condition | Coverage A (%) | Coverage A+B (%) | Coverage C+D (%) | A enrichment factor | A+B enrichment factor | C+D enrichment factor |
| --- | --- | --- | --- | --- | --- | --- |
| Baseline (30th / 70th pct) | 10.06 | 33.04 | 66.96 | 1.596 | 1.417 | 0.794 |
| Consensus 20th pct | 6.97 | 22.43 | 77.57 | 1.858 | 1.620 | 0.821 |
| Consensus 40th pct | 12.71 | 42.26 | 57.74 | 1.439 | 1.297 | 0.783 |
| Coherence 60th pct | 13.53 | 33.04 | 66.96 | 1.531 | 1.417 | 0.794 |
| Coherence 80th pct | 7.04 | 33.04 | 66.96 | 1.619 | 1.417 | 0.794 |

**Table S9.** Logistic regression of Scenario D and Scenario A membership on repeat-derived sequence content (>50% of window), adjusted for applicability-domain distance (standardized to one SD) and GC content, on the chr8/chr9 held-out test set, the full 517,790-window dataset (held-out test set and training pool combined), and the training pool alone. Odds ratios (OR) and 95% confidence intervals are shown for the repeat-derived, applicability-domain-distance, and GC-content terms of each model; full coefficient tables, including the intercept and (for the full dataset) the holdout-membership term, are in results/repeat_blindspot_results.json. Referenced from the main text’s Results and Discussion, Section 3.4.

| Dataset | n | Outcome | Repeat-derived OR (95% CI) | AD distance OR (95% CI) | GC OR (95% CI) |
| --- | --- | --- | --- | --- | --- |
| Held-out test (chr8/chr9) | 41,375 | Scenario D | 0.78 (0.75–0.81) | 1.43 (1.40–1.46) | 1.39 (1.10–1.75) |
| Held-out test (chr8/chr9) | 41,375 | Scenario A | 1.39 (1.30–1.49) | 0.59 (0.56–0.61) | 0.24 (0.17–0.35) |
| Full dataset | 502,838 | Scenario D | 0.79 (0.78–0.80) | 1.41 (1.40–1.42) | 1.28 (1.20–1.37) |
| Full dataset | 502,838 | Scenario A | 1.39 (1.37–1.42) | 0.61 (0.60–0.61) | 0.35 (0.31–0.39) |
| Training pool only | 461,463 | Scenario D | 0.79 (0.78–0.80) | 1.41 (1.40–1.41) | 1.27 (1.18–1.36) |
| Training pool only | 461,463 | Scenario A | 1.39 (1.36–1.42) | 0.61 (0.60–0.62) | 0.36 (0.32–0.41) |

### Supplementary Note: Comparison of bin-overlap and motif-span causal coherence

Causal coherence showed a non-monotonic relationship with absolute error across deciles. Genome-wide mean absolute error decreased from 0.503 in the lowest causal-coherence decile to a minimum of 0.299 in decile 8, then increased to 0.499 in decile 10. This extreme-tail association was tested against six candidate explanatory variables, target magnitude, applicability-domain distance, motif-instance count, GC content, repeat-derived content, and the instance density of the single most enriched panel transcription factor in the tail (KLF1), each of which explained only a small part of the association, and a small residual association persisted after adjusting for all six jointly (partial rho ~ +0.05, genome-wide). The effect was not statistically distinguishable from zero within the held-out test set alone and therefore was not independently confirmed in the reliability-scoped subset; the smaller sample size provides substantially less power to detect an effect of the genome-wide magnitude. This tail behaviour is therefore reported as incompletely explained rather than attributed to any single cause, and was not used to define the operational trust taxonomy.

**Table S10.** Full-dataset A/B/C/D scenario coverage under the bin-overlap (operational) and motif-span causal coherence definitions. The causal definition uses its own separately calibrated agreement cutoff (0.2218), reused unchanged from the held-out calibration; “Held-out crosscheck” reproduces the taxonomy on the chr8/chr9 subset within the full-dataset run as a consistency check against the primary held-out-only run, which differs by at most one window per class (Table 1 for the bin-overlap primary held-out run). Referenced from the main text's Section 3.7.

| Definition | Population | n (with shell) | A n (%) | B n (%) | C n (%) | D n (%) |
| --- | --- | --- | --- | --- | --- | --- |
| Bin-overlap | Full dataset | 502,898 | 49,894 (9.92%) | 114,691 (22.81%) | 91,948 (18.28%) | 246,365 (48.99%) |
| Bin-overlap | Held-out crosscheck | 41,379 | 4,161 (10.06%) | 9,511 (22.99%) | 7,347 (17.76%) | 20,360 (49.20%) |
| Causal | Full dataset | 502,898 | 63,306 (12.59%) | 101,278 (20.14%) | 88,657 (17.63%) | 249,657 (49.64%) |
| Causal | Held-out crosscheck | 41,379 | 5,214 (12.60%) | 8,458 (20.44%) | 7,069 (17.08%) | 20,638 (49.88%) |

**Table S11.** Matched-sample-size comparison of bin-overlap and motif-span causal coherence, held-out test set, at the operational Scenario A sample size (n = 4,161 highest-coherence windows within the high-consensus population, versus the remaining n = 9,511 high-consensus windows). Referenced from the main text's Section 3.7.

| Definition | Group A mean\|error\| | Group B mean\|error\| | Mann-Whitney p (A<B) | Welch t p | Precision ratio A/B @0.1 | @0.2 | @0.3 |
| --- | --- | --- | --- | --- | --- | --- | --- |
| Bin-overlap | 0.3959 | 0.4349 | 9.6e-10 | 3.2e-09 | 1.192 | 1.130 | 1.091 |
| Causal | 0.3816 | 0.4411 | <1e-300 | 4.6e-18 | 1.692 | 1.519 | 1.348 |

**Table S12.** Causal-coherence decile profile: mean absolute error by causal-coherence decile (decile 1 = lowest causal coherence), held-out test set and full dataset. Referenced from the main text's Section 3.7 and the Supplementary Note above.

| Decile | Held-out n | Held-out mean\|error\| | Full-dataset n | Full-dataset mean\|error\| |
| --- | --- | --- | --- | --- |
| 1 | 1,367-1,368 | 0.5241 | 16,458-16,459 | 0.5029 |
| 2 | 1,367-1,368 | 0.5441 | 16,458-16,459 | 0.4833 |
| 3 | 1,367-1,368 | 0.4907 | 16,458-16,459 | 0.4488 |
| 4 | 1,367-1,368 | 0.4270 | 16,458-16,459 | 0.4069 |
| 5 | 1,367-1,368 | 0.3957 | 16,458-16,459 | 0.3766 |
| 6 | 1,367-1,368 | 0.3642 | 16,458-16,459 | 0.3380 |
| 7 | 1,367-1,368 | 0.3401 | 16,458-16,459 | 0.3154 |
| 8 | 1,367-1,368 | 0.3070 | 16,458-16,459 | 0.2986 |
| 9 | 1,367-1,368 | 0.3183 | 16,458-16,459 | 0.3129 |
| 10 | 1,367-1,368 | 0.5190 | 16,458-16,459 | 0.4992 |

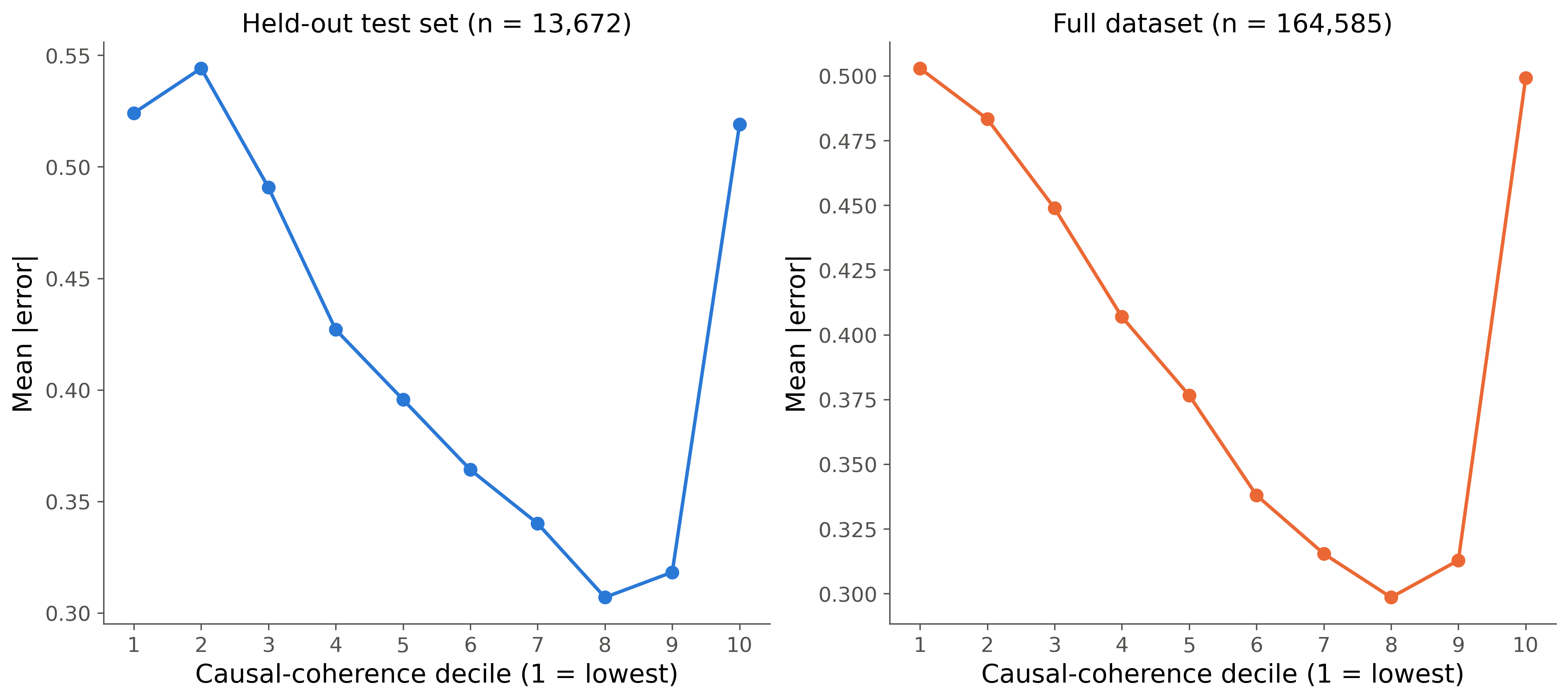

**Figure S3.** Causal coherence shows a non-monotonic (U-shaped) relationship with prediction error across deciles. Mean absolute error by causal-coherence decile (1 = lowest coherence, 10 = highest), for (A) the held-out chr8/chr9 test set and (B) the full 517,790-window dataset. Both curves fall to a minimum at decile 8 and rise sharply in the extreme top decile, the residual signal examined in the confound-controlled analyses of Table S13.

**Table S13.** Confound-controlled partial Spearman correlations between coherence and absolute error, across increasing control sets (3-control: |y_true|, applicability-domain distance, motif-instance count; 5-control: adds GC content and repeat-derived fraction; 6-control: adds KLF1 instance density). Population sample sizes for the held-out rows vary by 1-6 windows across the three control-set scripts (13,669-13,672 for whole A+B; 1,366-1,367 for the top decile) because of small differences in row exclusion between scripts; this does not affect the qualitative pattern. Referenced from the main text's Section 3.7 and the Supplementary Note above.

| Coherence | Population | Dataset | n | Raw $\boldsymbol{\rho}$ | 3-control partial $\boldsymbol{\rho}$ (p) | 5-control partial $\boldsymbol{\rho}$ (p) | 6-control partial $\boldsymbol{\rho}$ (p) |
| --- | --- | --- | --- | --- | --- | --- | --- |
| Causal | Whole A+B | Full dataset | 164,585 | -0.1702 | -0.0423 (5.9e-66) | -0.0505 (3.0e-93) | -0.0507 (6.2e-94) |
| Causal | Whole A+B | Held-out only | 13,669-13,672 | -0.1810 | -0.0292 (6.3e-04) | -0.0413 (1.8e-06) | -0.0408 (1.8e-06) |
| Causal | Top decile | Full dataset | 16,458 | 0.1437 | 0.0669 (8.3e-18) | 0.0570 (2.5e-13) | 0.0535 (6.6e-12) |
| Causal | Top decile | Held-out only | 1,366-1,367 | 0.1404-0.1413 | 0.0105 (0.697) | 0.0035 (0.898) | -0.0040 (0.883) |
| Bin-overlap | Whole A+B | Full dataset | 164,585 | -0.0476 | -0.0277 (3.2e-29) | -0.0238 (4.5e-22) | -0.0216 (1.9e-18) |
| Bin-overlap | Whole A+B | Held-out only | 13,669-13,672 | -0.0569 to -0.0572 | -0.0365 (2.0e-05) | -0.0297 (5.2e-04) | -0.0265 (1.9e-03) |
| Bin-overlap | Top decile | Full dataset | 16,458 | -0.0503 | -0.0119 (0.126) | -0.0136 (0.082) | -0.0124 (0.111) |
| Bin-overlap | Top decile | Held-out only | 1,366-1,367 | -0.0803 to -0.0817 | -0.0654 (0.0156) | -0.0625 (0.0210) | -0.0638 (0.0183) |

**Table S14.** Coherence-error correlation and A/B/C/D scenario composition, in-domain (ID) versus out-of-domain (OOD) at the calibrated applicability-domain cutoff, held-out test set and full dataset, both coherence definitions. Referenced from the main text's Section 3.8.

| Coherence | Dataset | Subset | n | coherence-error $\boldsymbol{\rho}$ (p) | A% | B% | C% | D% |
| --- | --- | --- | --- | --- | --- | --- | --- | --- |
| Bin-overlap | Held-out | ID | 39,293 | -0.0385 (2.3e-14) | 10.32 | 23.57 | 17.55 | 48.56 |
| Bin-overlap | Held-out | OOD | 2,086 | 0.0113 (0.607) | 5.03 | 12.03 | 21.67 | 61.27 |
| Bin-overlap | Full dataset | ID | 476,762 | -0.0314 (5.2e-104) | 10.19 | 23.35 | 18.13 | 48.33 |
| Bin-overlap | Full dataset | OOD | 26,136 | -0.0079 (0.202) | 4.95 | 12.92 | 21.07 | 61.06 |
| Causal | Held-out | ID | 39,293 | -0.1318 (7.5e-152) | - | - | - | - |
| Causal | Held-out | OOD | 2,086 | -0.0086 (0.696) | - | - | - | - |
| Causal | Full dataset | ID | 476,762 | -0.1188 (<1e-300) | - | - | - | - |
| Causal | Full dataset | OOD | 26,136 | 0.0095 (0.125) | - | - | - | - |

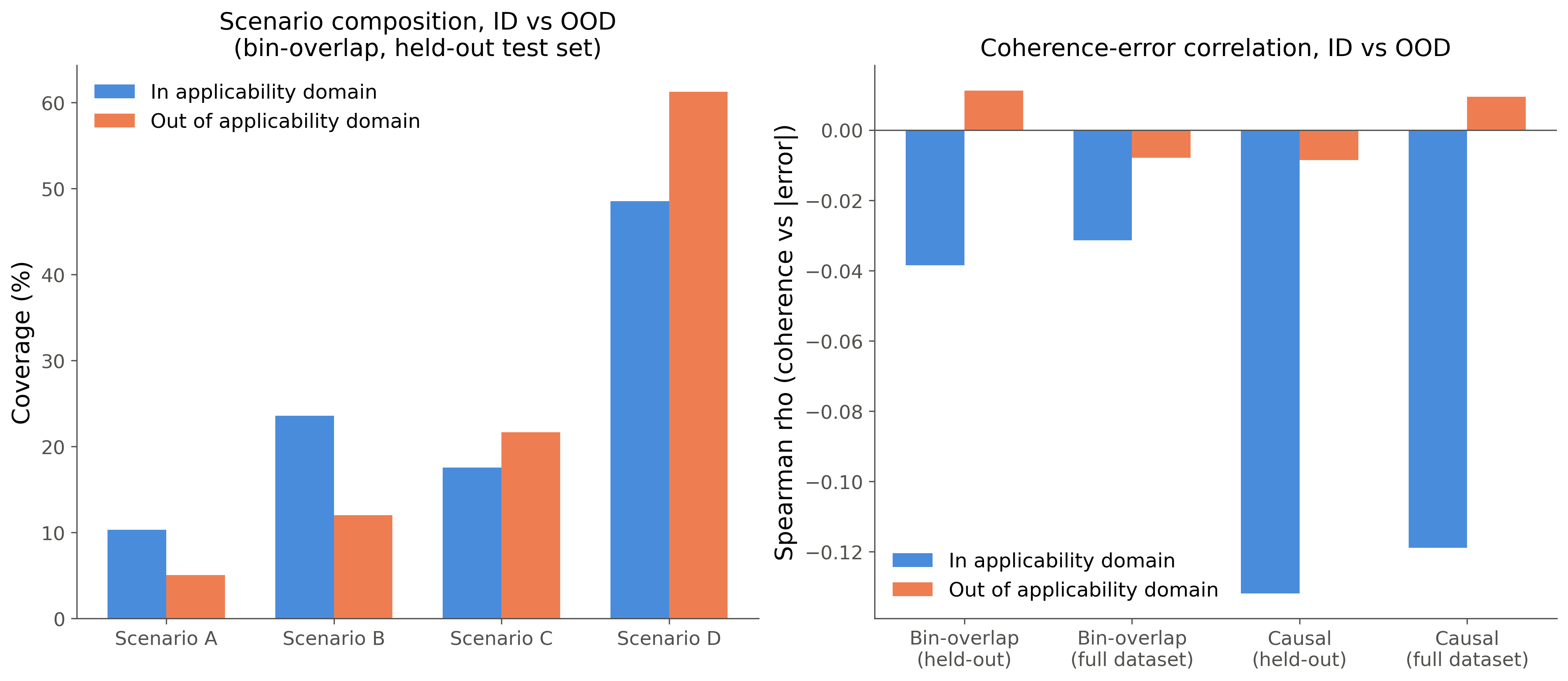

**Figure S4.** Explanation coherence provides little additional error discrimination outside the applicability domain, a population already depleted of high-consensus predictions. (A) Scenario A/B/C/D composition, in-domain (ID) versus out-of-domain (OOD), under the bin-overlap definition (held-out test set). OOD windows are disproportionately Scenario C/D. (B) Spearman correlation between coherence and absolute error, ID versus OOD, for both coherence definitions and both datasets; the ID-OOD gap narrows or reverses under the OOD selection effect described in the main text.

**Table S15.** Per-panel-transcription-factor instance counts, top causal-coherence decile versus the rest of the high-consensus (A+B) population, full dataset (n = 16,458 top decile, n = 148,127 rest). KLF1 is the only panel TF enriched, rather than depleted, in the top decile. Referenced from the Supplementary Note above.

| Panel TF | Top-decile mean instances | Rest-of-A+B mean instances | Ratio (top/rest) |
| --- | --- | --- | --- |
| GATA1 | 2.602 | 5.144 | 0.506 |
| GATA1::TAL1 | 2.214 | 4.634 | 0.478 |
| TAL1::TCF3 | 3.170 | 3.967 | 0.799 |
| KLF1 | 17.604 | 4.896 | 3.596 |
| NFE2 | 2.147 | 2.854 | 0.752 |
| MAF::NFE2 | 3.396 | 4.464 | 0.761 |
| GATA2 | 2.332 | 4.628 | 0.504 |
| RUNX1 | 6.357 | 6.424 | 0.990 |
| MYB | 2.464 | 2.105 | 1.171 |
| STAT5A::STAT5B | 6.338 | 7.973 | 0.795 |

**Table S16.** Superadditivity of joint versus summed motif-knockout effects for multi-instance motif modules, held-out test set and full dataset. Referenced from the main text's Section 3.7.

| Dataset | n multi-instance modules | Fraction superadditive | Fraction subadditive | Wilcoxon p (joint vs summed) |
| --- | --- | --- | --- | --- |
| Held-out test set | 387,835 | 91.07% | 8.93% | <1e-300 |
| Full dataset | 4,703,837 | 90.40% | 9.60% | <1e-300 |
